# Cellulose synthesis in Arabidopsis by heterologous expression of lower plants CESA proteins

**DOI:** 10.1101/2024.07.16.603775

**Authors:** Manoj Kumar, Leonardo D. Gomez, Simon Turner

**Affiliations:** University of Manchester; Faculty of Biology, Medicine and Health; Michael Smith Building; Oxford Road, Manchester M13 9PT, UK; Centre of Novel Agricultural Products (CNAP), Department of Biology, University of York, Heslington, York YO10 5DD, UK

## Abstract

Cellulose is the most abundant component of plant cell walls where it plays a pivotal role in regulating plant cell size and shape. In addition, as a component of the woody secondary cell walls, cellulose represents an abundant renewable resource to produce materials and chemicals. In higher plants, cellulose is synthesised at the plasma membrane by a hexameric protein complex, known as the rosette, that is able to synthesise 18 glucose chains that bond together to form a microfibril. While this rosette structure is highly conserved, significant variation exists in the structure and physical properties of cellulose found in different cell types and synthesised by different species. In this study, we surveyed the ability of the catalytic subunits of the cellulose synthase complex (CESA proteins) from a range of lower plant species to synthesise cellulose in the Arabidopsis secondary cell walls. Several lower plant CESA proteins are able to function in higher plants in conjunction Arabidopsis CESAs. Additionally, two moss CESA proteins synthesised cellulose in absence of Arabidopsis CESAs but with reduced crystallinity, indicating that it is the structure of CESA proteins themselves and not the cellular environment that determines the properties of the cellulose synthesised.

## Introduction

Global warming caused by anthropogenic emissions of greenhouse gasses has generated a dramatic increase in demand for materials and chemicals from renewable resources (de Vries *et al*., 2021). Plant biomass is an abundant renewable feedstock with the potential to satisfy this demand, making it one of the few resources able to make a meaningful contribution towards reducing global CO_2_ emissions. Consequently, there has been considerable investment in generating biofuels and chemicals using plant cell walls as a feedstock (de Vries et al., 2021). Recent progress in generating novel forms of engineered wood that includes flexible and mouldable wood (Xiao *et al*., 2021), densified wood (Song *et al*., 2018) and even transparent wood (Jia *et al*., 2019) have dramatically improved the mechanical properties of plant biomass, allowing it to be used in a much wider range of applications. Cellulose is the major constituent of plant biomass and is composed of linear chains of β1,4 glucose that associate into cellulose microfibrils to confer remarkable structural properties to this polymer. A variety of materials are generated from the breaking down of cellulose microfibrils, including micro-fibrillated cellulose and cellulose nanocrystals (Schubert *et al*., 2022, Ding *et al*., 2022). The properties of these novel products depend upon the properties of the cellulose-hemicellulose matrix of the plant cell wall, and additional improvements in the material properties could be attained if we were able to generate a variety of cellulose microfibrils *in planta*, with clearly different structural characteristics. In a similar way, modified cellulose in plants can modulate the recalcitrance to produce chemicals derived from plant biomass.

The cellulose microfibrils are the major load bearing component of plant cell walls. Consequently, these cellulose microfibrils have a major role in regulating cell expansion and plant growth. In plants, the catalytic subunits of the cellulose synthase complex (CSC), the protein complex responsible for the polymerisation of UDP-glucose into cellulose, are commonly known as the CESA proteins. In land plants (embryophytes) the CESA proteins form an 18mer that constitutes (Cosgrove, 2024, Purushotham *et al*., 2020, Fernandes *et al*., 2011) a hexagonal complex, known as a rosette, that is able to synthesise a cellulose microfibril as it moves through the plane of the plasma membrane. Apart from embryophytes, cellulose is produced by a diverse group of organisms including green algae (charophytes and chlorophytes), rhodophytes (red algae), xanthophytes (yellow-green algae), phaeophytes (brown algae), oomycetes, dinoflagellates, slime moulds, tunicates and bacteria (Niklas, 2004). There is wide variation in both the arrangements and number of the catalytic subunits within the complex synthesising cellulose and this in turn generates huge variation in the structure of the cellulose microfibril synthesised (Tsekos, 1999). This variation represents an as yet untapped source of structurally different cellulose with the potential for a variety of new biomaterials with improved chemical and physical properties.

Research in this area is hampered by a lack of basic information regarding the relationship between the structure of CESA proteins, their organisation within the cellulose synthase complex and the properties of the cellulose microfibril produced. In higher plants two distinct forms of the CSC form cellulose in either the primary or secondary cell wall (Cosgrove, 2024, Kumar & Turner, 2015a) that generate cellulose that varies in its physical properties, with cellulose from secondary cell wall found to have fewer amorphous regions than that found in primary cell walls. It remains unclear whether this variation is a consequence of the local environment in which the cellulose is synthesised or an inherent property of the complex that synthesises the cellulose.

In this report, we undertook a comprehensive study by expressing 15 different lower plant CESAs in Arabidopsis. We tested if they are functionally equivalent to those of Arabidopsis CESAs and whether they can function independently to synthesise cellulose in higher plants in the absence of any of the Arabidopsis secondary cell wall CESA proteins. We demonstrate that it is the CESA proteins *per se* and not the cell wall environment that determine cellulose properties. Furthermore, we demonstrate that by synthesising cellulose using heterologous expression of genes from other species, we are able to generate variation in the properties of the cellulose produced.

## Results

### Phylogeny of CESA proteins

We analysed a large number of manually curated CESA and CSLD sequences from across the taxonomic diversity of Streptophyta. A total of 6580 sequences plant CESA and CSLD sequences that possessed the typical features of rosette type CESA proteins were aligned and used to generate a phylogenetic tree (Figure 1 and S1). There was clear separation of CESA and CSLD sequences. CESA sequences from higher plants formed 6 separate clades for the 6 well established CESA phylogenetic classes named after the Arabidopsis members CESA1, CESA3, and CESA6 that are involved in primary cell wall biosynthesis and CESA4, CESA7 and CESA8 required for secondary cell wall synthesis (Figure 1 and S1). We also identified 3 CESA phylogenetic classes in ferns, Fern A, Fern B and Fern C, positioned with CESA1/3, CESA6 and CESA4/7/8 classes from angiosperms respectively. These results are comparable to those obtained by Pancaldi *et al*. (2022), the most comprehensive phylogenetic analysis of CESA sequences from seed plants published to date.

**Figure 1.**
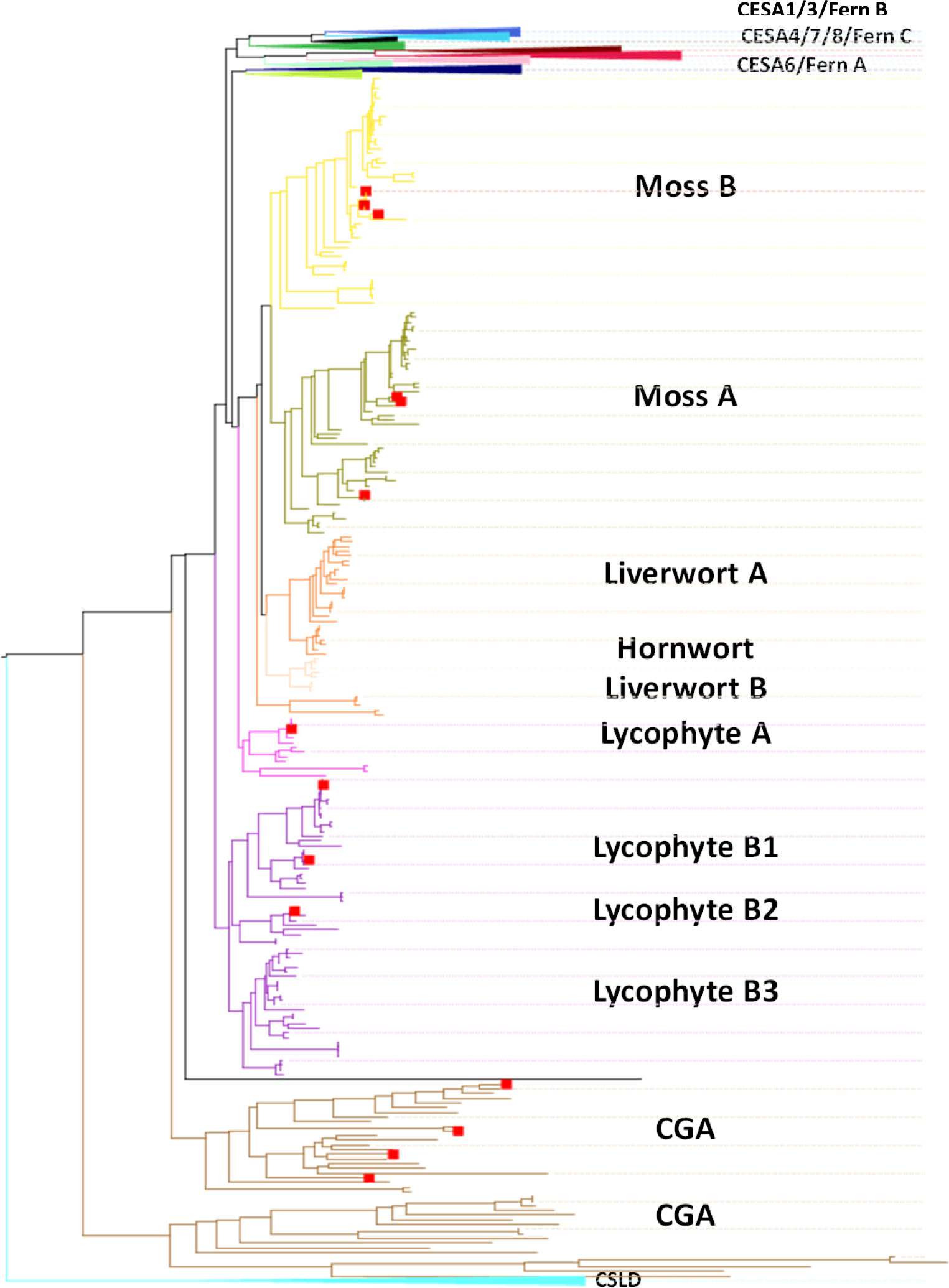
Phylogenetic tree showing classification of Streptophyta CESA and CSLD sequences. A selection of 6580 Streptophyta sequences containing the GT2 domain and the RING domain in their N-terminus were aligned with MUSCLE and a maximum likelihood (ML) tree was inferred using Fast-tree. Six phylogenetic classes of CESA proteins from seed plants are labelled based on their Arabidopsis members. The tree is organised to highlight the details of bryophyte, lycophyte and CGA sequences. 14 proteins used for complementation studies are indicated with red dots. An alternative representation of the same tree is shown in figure S1.

The CESA sequences from seed plants separate into distinct phylogenetic classes that include examples from dicots, monocots, gymnosperms within each class (Figure 1). The same is also true for fern sequences that also form three separate clades. In contrast the lower plant CESA sequences from bryophytes, lycophytes and CGA species do not mix with sequences from seed plants and ferns or each other, forming separate clades for each group (Figure 1, S1). However, when there are multiple CESA sequences present in these lower plant species, they do separate into “phylogenetic classes”. There appeared to be at least 2 distinct classes for each of these taxonomic groups (class A and class B). For lycophytes, class B is split into three further sub-groups. (Figure 1). However, the functional distinctness is only established for the moss *Physcomitrella patens* that has two distinct CESA classes with a member from each class being required for cellulose synthesis (Norris *et al*., 2017a, Scavuzzo-Duggan *et al*., 2018, Li *et al*., 2019). Within a phylogenetic tree of CESA proteins from green plants, these moss CESA proteins form a completely separate clade from higher plants CESAs (Figure 1).

### Functional analysis of lower plant CESA proteins

In order to test whether CESA proteins from lower plant species were functionally equivalent to Arabidopsis secondary cell wall CESA proteins, we expressed 15 selected lower plants CESAs into Arabidopsis *cesa*4, cesa7 and *cesa8* single and higher order mutants. The selected proteins covered a range of taxonomic diversity (Figure 1) and included one CESA protein from each of 4 different CGA species, 6 CESA proteins from the moss *Physcomitrella patens,* all 4 CESA proteins from spike moss *Selaginella moellendorffii* and a single CESA protein from the fern *Osmunda* that came from group C, the clade that appears to share a common origin with the secondary cell wall CESA proteins (Figure 1, S1). Despite the evolutionary distance the overall structure and large parts of the proteins are well conserved (Figure S2).

Genes corresponding to the selected 15 proteins, driven by Arabidopsis CESA7 protomer, were transformed into each Arabidopsis single secondary cell wall mutant, *ces4, cesa7,* or *cesa8*, as well as all double mutant combinations and the triple mutant leading to generation of nearly 100 genotypes. For complementation analysis, rosette area, plant height and cellulose content were measured. All genotypes were analysed at T1 stage while a selection of genotypes were also analysed again in the T2 generation. As a result of the large number of genotypes involved, plants were grown across multiple batches with the wild type and *cesa* mutant controls included with each batch (Figure S3). In order to facilitate comparisons between different batches, all data is expressed as % complementation (Kumar *et al*., 2018b). In this calculation, WT is 100% and the *cesa* mutant is 0%. We measured both cellulose content and plant height as we have previously observed that there is strong correlation between these two traits, but in some instances small increases in cellulose content lead to larger increases in plant height (Kumar et al., 2018b).

None of the CESA sequences from the 4 CGA species were able to complement the plant height, or cellulose content defects of the mutants (Figure 2A and S4A). These results indicate that these CESA proteins are unable to form functional complexes with either of the Arabidopsis *cesa4, cesa7* or *cesa8* mutants or form functional homomeric complexes in higher plants.

**Figure 2.**
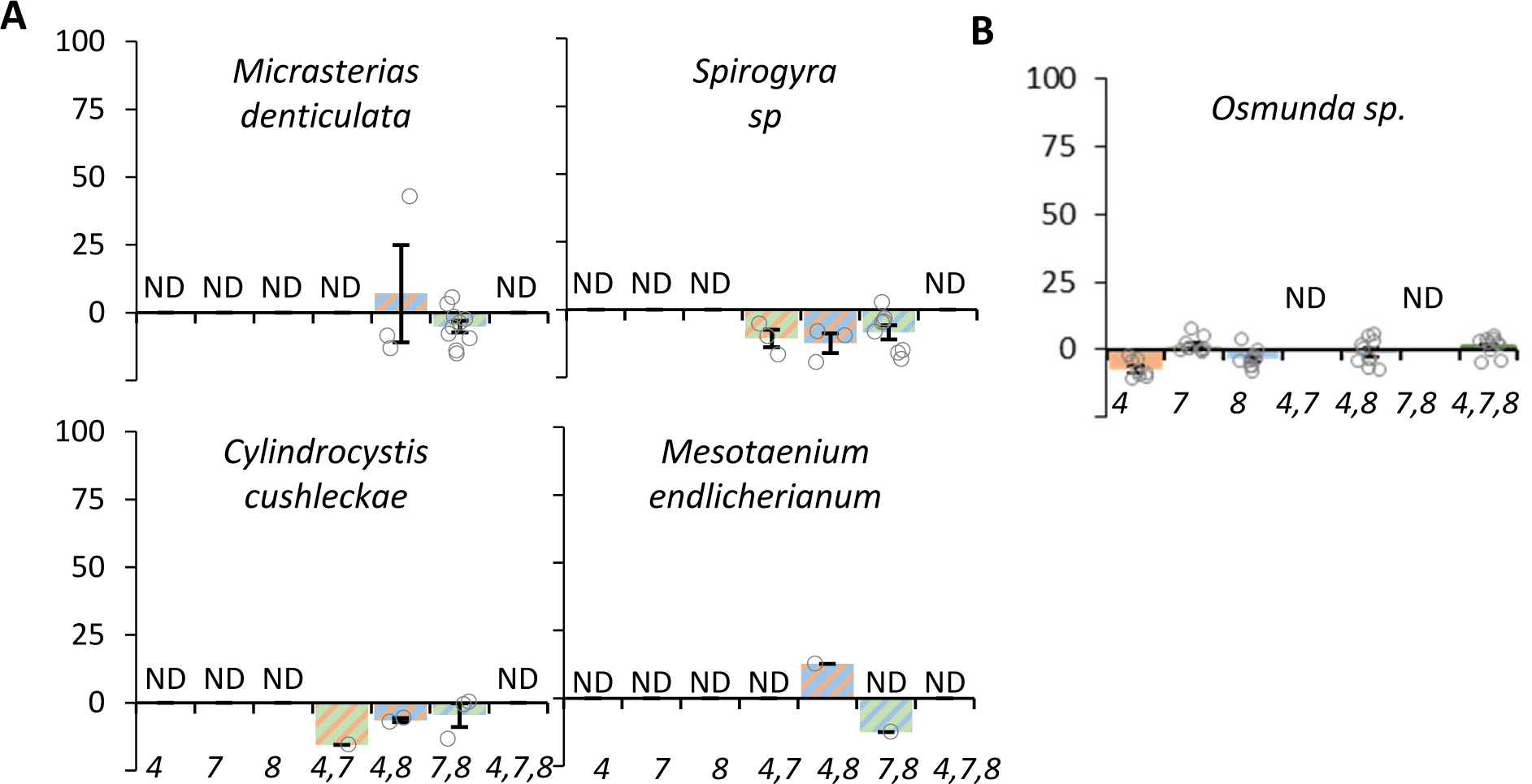
Complementation of the cellulose deficient phenotype of Arabidopsis secondary cell wall CESA mutants expressing CESA proteins from species of CGA and the fern *Osmunda*. Genes encoding single CESA proteins were expressed in Arabidopsis *cesa4,7,8* single and multiple mutant knock-out combination. (A) Genes encoding CESA proteins from 4 species of CGAs. (B) A gene encoding a CESA proteins from fern *Osmunda species,* a member of fern group C. Result are expressed as % complementation. *Cesa* mutants are indicated by the number in italics on the x axis. Up to 10 individual T1 plants were measured for each genotype and mean values are shown with standard error bars. Individual measurements are indicated by open circles. No significant complementation was observed and judged by univariate ANOVA between the transgenic lines and the *cesa4,7,8* triple mutant. ND – not determined.

Since fern class C appears related to higher plant secondary cell wall CESA proteins but appears to have arisen prior to the diversification of the secondary cell wall CESA proteins into different classes (Figure 1, S1), we hypothesised that this fern group might form homomeric rosettes. To test this hypothesis, we analysed the ability of a class C CESA proteins from the fern *Osmunda* to complement the cesa4, cesa7, and *cesa8* mutants. As with the CGA genes, we observed no complementation using this fern gene (Figure 2B and S4B).

Genetic analysis of the moss *Physcomitrella patens* has previously demonstrated that CESAs genes fall into two distinct phylogenetic and functional classes (Norris et al., 2017a, Tran *et al*., 2018, Li et al., 2019). Class moss A includes PpCESA3, 5 and 8 while moss B includes PpCESA4, 6, 7 and 10 (Scavuzzo-Duggan et al., 2018). CESAs from both classes are involved in cellulose synthesis in primary cell walls (PpCESA5, 4 and 10) and secondary cell walls (PpCESA3, 8, 6 and 7) (Norris et al., 2017a, Li et al., 2019). Among the moss A CESAs, PpCESA8 was able to partially, but significantly, complement both the cellulose deficient and plant height defect of the *cesa4^irx5-4^* mutant (Figure 3A, S5A). Additionally, some individual complemented T1 lines were obtained when PpCESA3 and 5 were expressed in *cesa4^irx5-4^* mutant, although the results were not statistically significant when tested across 10 independent T1 lines (Figure 3A). Moss A CESAs did not complement any of the other single, double or triple *cesa* mutants. Since we observed no complementation of the double mutant combinations or the triple mutants the results suggest that members of the moss class A CESA proteins are able to function together with the Arabidopsis CESA7 and CESA8 proteins to form a functional complex.

**Figure 3.**
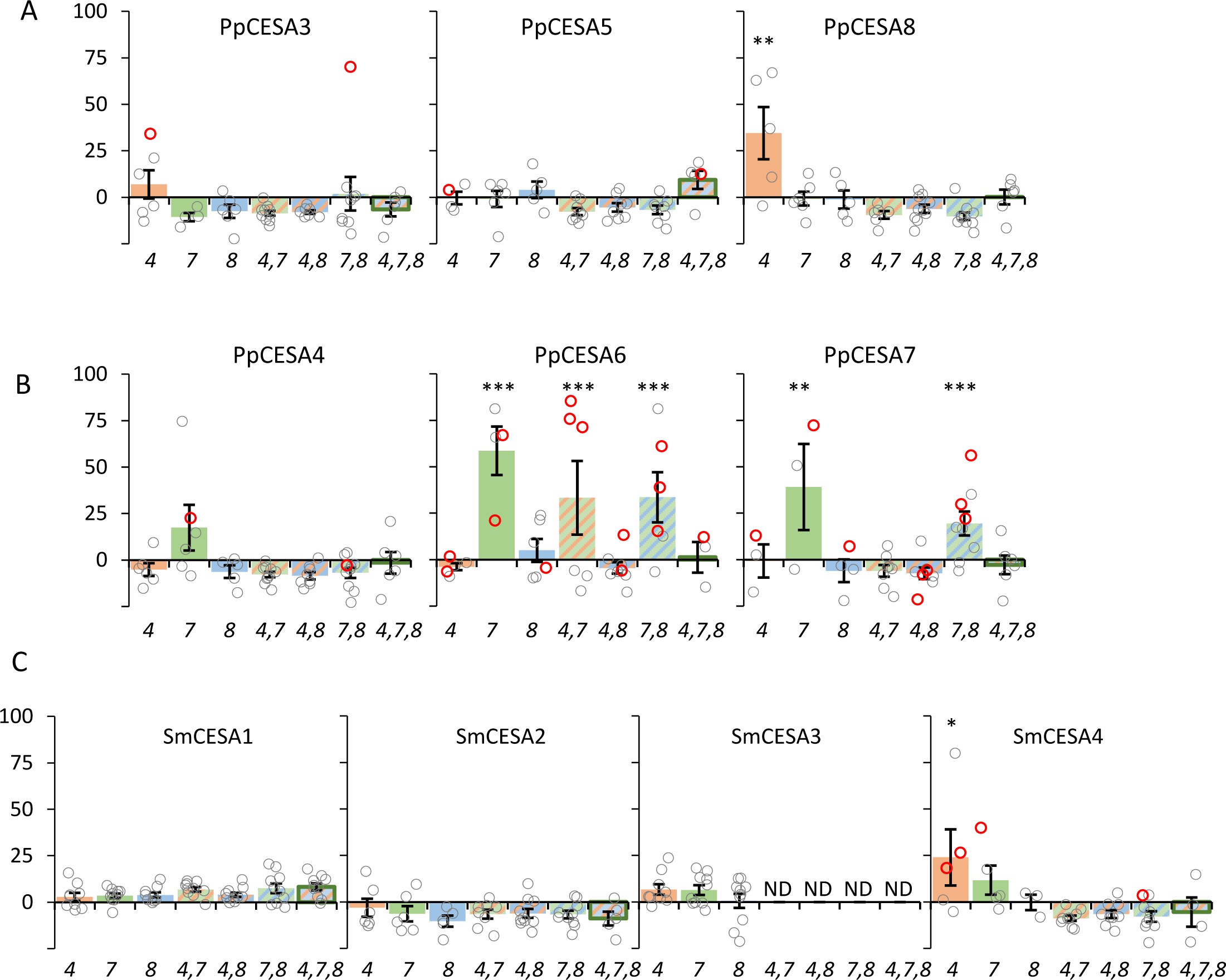
Complementation of the cellulose deficient phenotype of Arabidopsis secondary cell wall *cesa* mutants expressing CESA proteins from the moss, *Physcomitrella patens* and the spikemoss, *Selaginella moellendorffii*. Individual CESA protein were expressed in Arabidopsis *cesa4,7,8* single and multiple mutant knock-out combination mutants. *Cesa* mutants are indicated by the number in italics on the x axis. Result are expressed as % complementation. (A) Genes encoding CESA proteins from moss, *Physcomitrella patens, moss group A*. (B) Genes encoding CESA proteins from moss, *Physcomitrella patens, moss group* B. (C) Genes encoding 4 different CESA proteins from *Selaginella moellendorffii*. Up to 10 individual T1 plants were measured for each genotype and mean values are shown with standard error bars. Empty circles represent all individual measurements and the lines selected for T2 analysis are indicated with red circle. Significantly positive complementation, calculated using univariate ANOVA between the genotype and the *cesa4,7,8* triple mutant, is indicated with asterisks. *** Significant at 0.001, ** significant at 0.01, * significant at 0.05. ND – not determined.

When we analysed complementation following expression of the moss B CESAs, PpCESA4, 6 and 7, we were able to obtain significant complementation of the cellulose content of the *cesa7^irx3-5^* mutant with both PpCESA6 and PpCESA7 proteins (Figure 3B and S5B) while all three class B Physcomitrella genes were able to complement the plant height phenotype of *cesa7^irx3-5^*. Furthermore, both PpCESA 6 and 7 were also able to complement *cesa8^irx1-7^* mutant albeit to a lesser extent than the *cesa7^irx3-7^* mutant, with only the complementation observed with PpCESA6 being statistically significant (Figure 3B). None of the moss B CESAs were able to complement *cesa4^irx5-4^* mutant. Moreover, PpCESA6 and PpCESA7 were both able to complement *cesa7,cesa8* double mutant and PpCESA6 also complemented the *cesa4,cesa7* double mutant. None of the individual Physcomitrella CESAs were able to complement the *cesa4 cesa7 cesa8* triple mutant (Figure 3B and S5B).

We also tested all 4 CESA proteins from Selaginella, a lycophyte, for their ability to complement Arabidopsis *cesa4*, *cesa7* and *cesa8* mutants (Figure 3C, and S5C). Out of the 4 CESAs genes, only SmCESA4 exhibited any significant complementation. It was able to complement both the cellulose content and plant height defects of *cesa4*. None of the Selaginella genes were able to complement any of the Arabidopsis *cesa* higher order mutants (Figure 3C and S5C).

In order to verify this data, we selected up to 3 independent lines from each genotype that exhibited at least some level of complementation and analysed the T2 generation. The results were broadly similar to those obtained in T1 generation with all three class A CESA genes from moss able to complement *cesa4* (Figure 4A and S6). Similarly, all three moss class B genes were able to complement the cesa7 mutant (Figure 4B and S5). Complementation with PpCESA6 was particularly effective as it was able to complement the *cesa4, cesa7,* and *cesa7,cesa8* double mutants in addition to *cesa7* and *cesa8* single mutants. Lines expressing SmCESA4 were able to complement both *cesa4* and *cesa7* mutants (Figure 4C).

**Figure 4.**
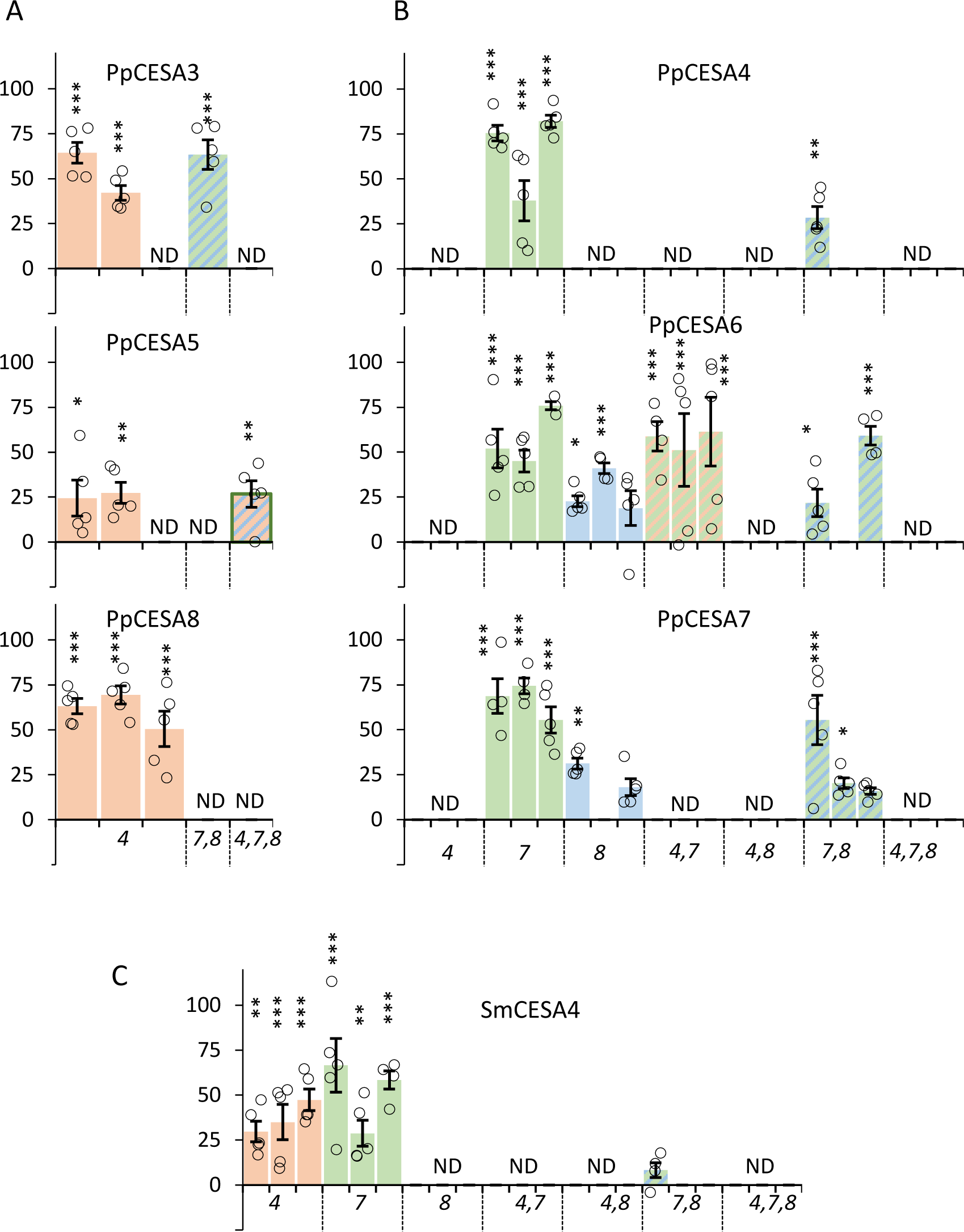
Complementation of **Cellulose content** phenotype of T2 generation of Arabidopsis SCW CESA mutants expressing CESA proteins from Physcomitrella class A (A), class B (B) and Selaginella (C). For selected genotypes, 3 independent T2 lines were grown and 5 plants were analysed for each line. Plant height data expressed as % complementation is shown. Empty circles represent individual measurements. Significantly positive complementation, calculated using univariate ANOVA between the genotype and the *cesa4,7,8* triple mutant, is indicated with asterisks. *** Significant at 0.001, ** significant at 0.01, * significant at 0.05.

### Moss CESA proteins alone are able to synthesises cellulose in higher plants

Our experiments with expression of single moss CESAs showed that class B moss CESAs, PpCESA6 and 7 were both able to complement *cesa7^irx3-7^* mutant and also *cesa8^irx1-7^*mutant to a lesser extent but not the *cesa4^irx5-4^* mutant. In contrast, PpCESA8, a class A moss CESA, effectively complemented the *cesa4^irx5-4^*mutant. This data is consistent with previous genetic analysis in moss which demonstrated that members from both class A and B were required to synthesise cellulose. With that in mind, we created two plasmids containing either both PpCESA6 and PpCESA8 or PpCESA7 and PpCESA8 and expressed them in the *cesa4,cesa7, cesa8* triple mutant background. Both combinations were able to significantly complement the triple mutant for plant height, rosette area and cellulose content phenotypes (Figure 5), indicating that the pairs of Physcomitrella genes were able to synthesis cellulose in the secondary cell wall independently of any Arabidopsis genes.

**Figure 5.**
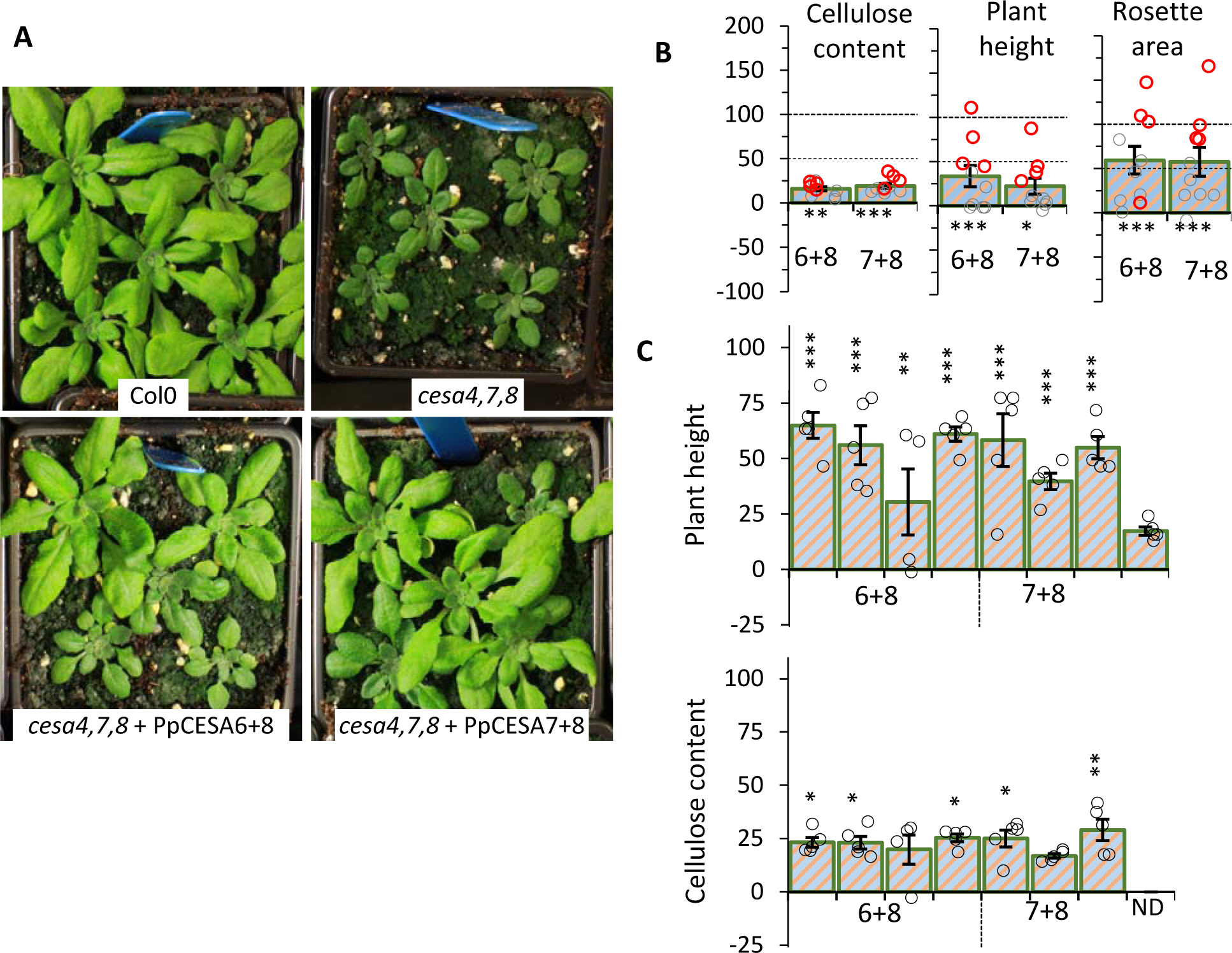
Complementation of Arabidopsis *cesa4,7,8* triple mutant by expressing two Physcomitrella CESA proteins. PpCESA6 and PpCESA8 (6+8) or PpCESA7 and PpCESA8 (7+8) were expressed in Arabidopsis *cesa4,7,8* triple mutant. (A) Images of representative plants at 4 weeks old stage. (B) Complementation of stem cellulose content, plant height and rosette area. 10 individual T1 plants were measured for each genotype. Empty circles represent all individual measurements and the lines selected for T2 analysis are indicated with red circles. (C) Complementsyion of cellulose content and plant height phenotype of selected T2 lines. Significantly positive complementation, calculated using univariate ANOVA between the genotype and the *cesa4,7,8* triple mutant, is indicated with asterisks. *** Significant at 0.001, ** significant at 0.01, * significant at 0.05.

While we observed significant complementation of cellulose content in these plants overexpressing 2 Physcomitrella CESA genes, we observed much larger changes in plant height and rosette area with some of the complemented lines growing as well as the wild type (Figure 5A, B). We have previously performed complementation analyses similar to the ones presented here and observed that the plant height phenotype is a more sensitive indicator of complementation where relatively smaller increases in cellulose content are able to lead to larger increases in plant height (Kumar *et al*., 2017, Kumar *et al*., 2018a). However, the discrepancy between cellulose content and plant height for the triple mutant complemented with two moss CESAs was greater than anything we have observed before or with the other genotypes presented in this study. To study this result in more detail, we selected 4 independent lines for each genotype and analysed the T2 generation. The results confirmed that the results obtained for the T1 plants, with all T2 lines exhibiting good complementation of plant height, but much smaller, albeit significant complementation of the defect in cellulose content (Figure 5C).

All our cellulose content measurements are performed using a variation of Updegraf’s method that only measures crystalline cellulose (Updegraff, 1969). Solid state nuclear magnetic resonance (ssNMR) can detect peaks for interior cellulose chains (indicative of crystalline cellulose) and surface cellulose chains (indicative of amorphous cellulose) (Dupree *et al*., 2015). To obtain a clearer overview of changes in cell wall composition of these lines, we used ssNMR to analyse Col0 WT, *cesa4,cesa7*,*cesa8* triple mutant, and 4 independent lines for *cesa4,cesa7*,*cesa8* triple mutant complemented with PpCESA6 and PpCESA8. As controls we also analysed genotypes from this study that exhibited good complementation for both plant height and cellulose content and included one line for each PpCESA6 transformed into the *cesa7* single and the cesa*4,cesa7* and *cesa7,cesa8* double mutants and a line with PpCESA8 transformed into the *cesa4* mutant (Figure 6). We paid particular attention to those peaks that are associated with interior and exterior cellulose chains including the pair of peaks at 74.5 and 74.2 ppm associated with C4 and those at 65.1 and 63ppm associated with C5 that correspond to the surface and interior chains respectively. Comparison on the signal from wild type samples with those from the *cesa4 cesa7 cesa8* triple mutant clearly shows big differences in peaks corresponding to cellulose including the 4 peaks listed above (Figure 6). For the control lines, the signal for both interior and exterior chains increased proportionately to generate a spectrum that more closely resembles the wild type (Figure 6). In contrast, for the lines expressing the Physcomitrella genes PpCESA6 and PpCESA8, there are a significant increase in the signal for exterior cellulose peaks at 74.5 and 65.1, but the signal from the interior chains at 74.2 and 63ppm was significantly smaller (Figure 6). This result indicates the presence of increased non-crystalline or amorphous cellulose in these lines.

**Figure 6.**
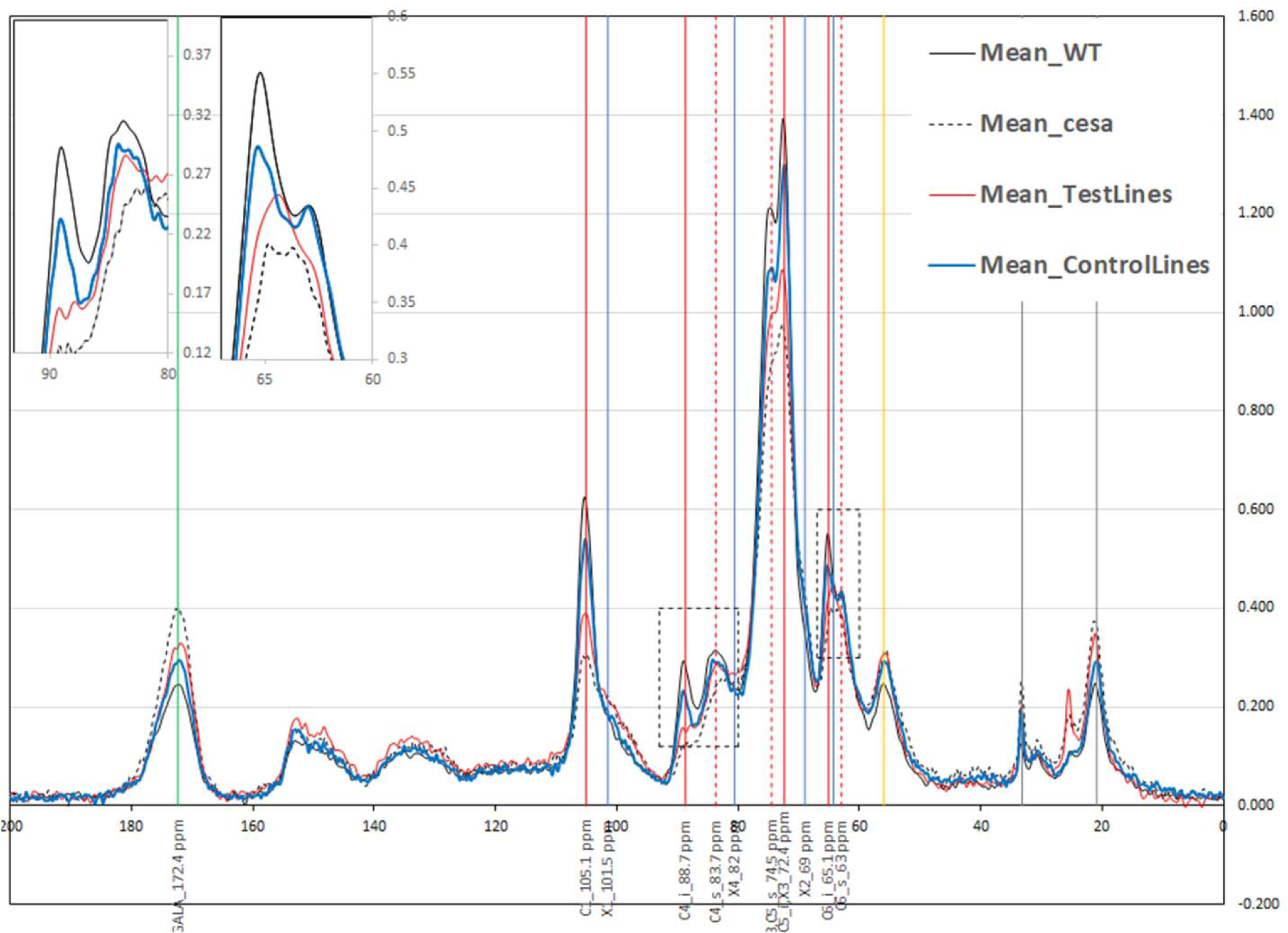
Solid state NMR analysis of Arabidopsis *cesa4,7,8* triple mutant complemented with Physcomitrella genes PpCESA6 and PpCESA8. Traces are the mean of spectra obtained from 4 different samples. Inset shows close up of regions associated with surface and interior cellulose sugars. “TestLines” is mean of 4 independent lines of *cesa4,7,8* triple mutant transformed with PpCESA6 and PpCESA8 while the “ControlLines” is mean of 4 different genotypes – PpCESA6 transformed into Arabidopsis *cesa7* mutant; PpCESA6 transformed into Arabidopsis *cesa4 cesa7* mutant; PpCESA6 transformed into Arabidopsis *cesa7 cesa8* mutant and PpCESA8 transformed into Arabidopsis *cesa4* mutant. Vertical lines indicate positions for peaks associated with cellulose (red lines - C4_i_88.7 ppm, C4_s_83.7 ppm, C6_i_65.1 ppm, C6_s_63 ppm, C1_105.1 ppm, C2,C5_i,X3_72.4 ppm, C3,C5_s_74.5 ppm); xylan (blue lines - X1_101.5 ppm, X2_69 ppm, X4_82 ppm, X5_64.3 ppm); aliphatic compounds (grey lines - ALP_20.9 ppm, ALP_33.2 ppm); carbonyl group (yellow line - Carbonyl_56 ppm); galacturonic acid (green line - GALA_172.4 ppm).

### Improved saccharification in lines synthesising cellulose co-expressing PpCESA6 and PpCESA8

To test whether the increased proportion of non-crystalline cellulose we observed in plants overexpressing 2 Physcomitrella genes might have any practical applications, we performed a digestibility assay on these samples. In a saccharification assay measuring sugar release from samples without any pretreatment, we observed a significant increase in sugar release using samples from 4 independent lines expressing both PtCESA6 and PtCESA8 in the *cesa4/cesa7/cesa8* triple mutant compared to either the wild type or the *cesa4,cesa7,cesa8* triple mutant controls (Figure 7). This would support the idea that the less crystalline cellulose generated by the PhCESA6 and PhCESA8 is more easily digested by cellulases. It is important to note that these plants grow normally and can achieve 100% plant height complementation in some cases (Figure 5B) meaning that there is little, or no growth penalty involved. These plants provide an example how cellulose synthesis could be manipulated to produce an improved feedstock for biobased chemicals and fuels.

**Figure 7.**
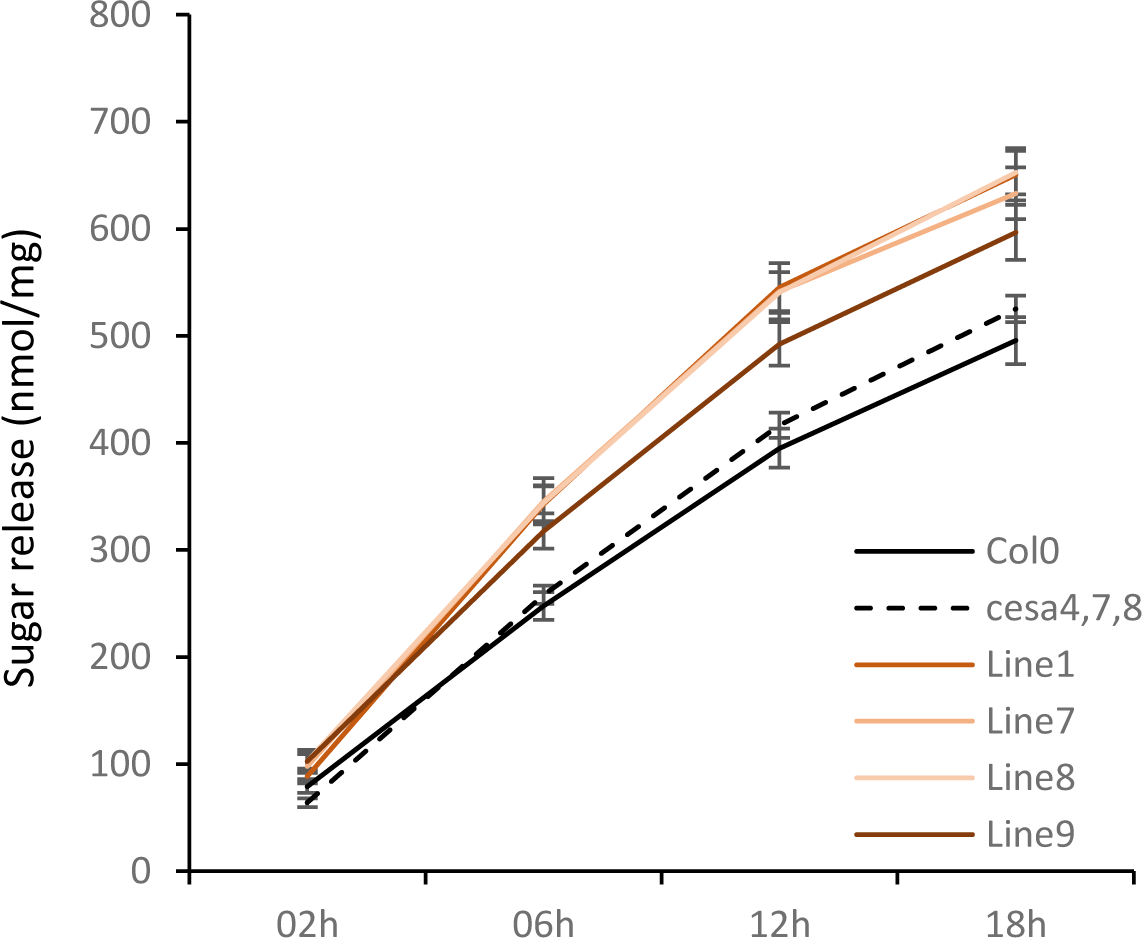
Digestibility assays of Arabidopsis *cesa4,7,8* triple mutant complemented with Physcomitrella genes PpCESA6 and PpCESA8. Sugar release data, without any pre-treatment for 4 independent lines of PpCESA6 and PpCESA8 transformed into *cesa4,7,8* triple mutant is shown. Each datapoint is a mean of 24 replicates (6 biological x 4 technical). Error bars are standard error of mean. Sugar release from all 4 lines significantly more at 6, 12 and 18h with p-value <0.05.

## Discussion

A previous study demonstrated partial complementation of the Arabidopsis primary cell wall cellulose mutant, *cesa6^prc^* using an atypical CESA protein from the marine red alga *Calliarthron tuberculosum* that contained an additional carbohydrate binding domain (CBD48), however it was unable to complement either the *cesa7*, mutant used in this study, or the c*esa3* mutants (Xue *et al*., 2022). Phylogenetic analysis suggests the CGA sequence show little evidence of gene duplication and diversification, and it is likely that *in situ* most CGA species are able to form homomeric rosette structures. Despite the conservation of their overall CESA structure and large regions of high sequence conservation, these CESAs from the CGAs appear to be sufficiently diverse from those sequences in higher plants that they are unable to synthesise cellulose in higher plants. While the proteins appear structurally conserved, it is clear that CESA proteins in higher plants require extensive post-translation modification (Kumar *et al*., 2016, Mergner *et al*., 2020, Chen *et al*., 2010) and the CESA proteins from CGA species may not be modified appropriately. Similarly, based upon the phylogenetic data, we hypothesise that fern class C CESA proteins are likely to form a homomeric complex, but the single CESA gene we tested from this group failed to complement *cesa* mutants, potentially because the proteins have diverged considerably from its common ancestor with the higher plant secondary cell wall CESA proteins (Figure S1). Among the 15 CESA sequences tested, all possess identical sequences at the 4 main catalytic motifs; DD (DDG), DxD (DCD), TED (TED) QxxRW (QVLRW) and the gating loop FxxVxK (FTVTS/AK) (Figure S2). Availability of crystal structure of a homo-trimer of heterologously expressed PttCESA8 revealed 25 contact points between different sub-units, located mainly in the P-CR, TM4 and TM7 regions (Purushotham et al., 2020). However, all these points are identical in nearly all Arabidopsis CESAs and the 15 lower plants CESAs tested here (Figure S2). Most differences among Arabidopsis and lower plant CESA sequences reside within and highly variable regions close to the N-terminus (VR1) and within the central cytoplasmic domain (VR2) (Figure S2). The structure of these regions is poorly understood and both contain known sites for post-translational modifications including phosphorylation (Mergner et al., 2020, Chen et al., 2010) and S-acylation (Kumar et al., 2016) as well as interacting with other proteins. As these regions are so variable between different CESA proteins, it is plausible that their failure to complement a particular *cesa* mutant, or to function independently during Arabidopsis secondary cell wall biosynthesis may result from either lack of or alternations in essential post-translational modifications or their inability to interact with other proteins essential for cellulose synthesis.

In contrast, the CESA genes from the moss Physcomitrella are clearly able to function in the higher plant Arabidopsis. It is notable that the two functional classes of CESA proteins identified in Physcomitrella are also required to synthesise cellulose in higher plants. While individual Physcomitrella CESA genes are unable to complement the *cesa4/cesa7/cesa8* triple mutant, a combination of the class A protein PpCESA8 and either of the class B genes PpCESA6 or PpCESA7 are able to complement the triple mutant. To our knowledge, this is the first report showing that *CESA* genes from higher plants can be completely replaced with that from a lower plant. The data also provides independent support of the studies in Physcomitrella that a CESA from both class A and B are required to make cellulose (Li et al., 2019, Norris *et al*., 2017b).

In our functional analysis of the Physcomitrella *CESA* genes, we find that *CESA* genes from moss class A are better at complementing Arabidopsis *cesa4* mutants, while those CESA genes from moss class B CESA are better at complementing *cesa7* and *cesa8* mutants (Figure 3, 4 S5 and S6). This is surprising, since based on phylogenetic analysis, CESA7 is believed to have initially split from the CESA4/CESA8 clade. Also, functional analyses of chimeric SCW CESA proteins involving domain swaps from other SCW CESAs, show that CESA7 exhibits little function complementation when domains from either CESA4 or CESA8 are swapped in, suggesting that CESA7 has particularly strict structural constraints (Kumar et al., 2017). The basis of this apparent promiscuity revealed for the structure of the moss class B CESAs is hard to understand in the absence of reliable structural data. While the structural information for the conserved catalytic domains of CESA proteins is available (Purushotham et al., 2020), the diverse regions likely to be responsible for the class specificity, such as the CSR region, are currently lacking. Once this information becomes available it should reveal important information on the fundamental rules governing CESA protein structure and assembly of the cellulose synthase complex.

Although crystallinity is one of the fundamental properties of cellulose, exactly what determines the proportion of ordered vs amorphous cellulose is unclear. Mutations in Arabidopsis CESA proteins are able to reduce the crystallinity of the cellulose synthesised (Harris *et al*., 2012), however this does not address what determines the differences in structural properties between the cellulose synthesised in primary cell walls compared to that synthesised in secondary cell walls or other examples of natural variation in cellulose structure. The cellulose synthesised by the moss CESA proteins in Arabidopsis was less crystalline than the cellulose synthesised in the secondary cell walls of the wild type (Figure 6). These alterations in the ratio of surface to interior cellulose are greater than those observed when we mutated the catalytic aspartates in CESA4, CESA7 or CESA8 (Kumar et al., 2018a), a change that we would anticipate would reduce the size of a microfibril by a third. This suggests that the changes observed in the cellulose produced by moss PpCESA8 with either PpCESA6 or PpCESA7 are not a consequence of simply altering the number chains within a microfibril. Our results support for the idea that the properties of cellulose are determined by the CESA proteins themselves and are not determined by the local environment in which the cellulose is synthesised.

One of the challenges of reducing greenhouse gas emissions is to use renewable materials. The sheer abundance of cellulose means it is one of the few renewable feedstocks with the potential to make a significant impact on reducing greenhouse gas emissions. The utility of cellulose as a feedstock could be increased by generating greater diversity of cellulose from abundant sources such as higher plants. For example, many red algae make a diverse array of much larger microfibrils (Tsekos, 1999) and if they could by synthesised in higher plants, it will offer huge potential for synthesising new material or acting as novel feedstocks. In the present work, we have demonstrated we are able to replace *CESA* genes from Arabidopsis with those from the moss Physcomitrella and generate low recalcitrance cellulose with altered properties that represents an important first step in generating new and improved materials and feedstocks from plant biomass.

## Methods

### Sequence retrieval

Three main strategies were used for building the collection of GT2 sequences, literature searches, BLAST searches and keyword searches. For literature search, sequences described in Blum *et al*. (2012) (oomycetes CESA sequences), Brawley *et al*. (2017), Chan *et al*. (2019), Little *et al*. (2018), Pang *et al*. (2020), Sena *et al*. (2019), Wang *et al*. (2020), Wan *et al*. (2018) were retrieved. For BLAST searches, 40 Arabidopsis GT2 sequences and a selection of 15 bacterial BcsA sequences were used as query to search fully sequenced genomes at phytozome (Goodstein *et al*., 2012). A local database was created to search fully sequenced genomes publicly available but not included in phytozome. This database included genomes from gymnosperms (*Cycas micholitzii, Ginkgo biloba, Gnetum montanum, Picea abies, Picea glauca, Pseudotsuga menziesii, Pinus pinaster, Picea sitchensis, Pinus sylvestris, Pinus taeda* all downloaded from https://bioinformatics.psb.ugent.be/plaza/versions/gymno-plaza/), ferns (*Azolla filiculoides* (Li *et al*., 2018)*, Salvinia cucullate* (Li et al., 2018)*, Ceratopteris richardii* (Marchant *et al*., 2019)*, Adiantum capillus-veneris* (Fang *et al*., 2021)), bryophytes (*Anthoceros angustus* (Zhang *et al*., 2020)*, Anthoceros agrestis Bonn, Anthoceros agrestis Oxford, Anthoceros punctatus* (Li *et al*., 2020)) and CGA (*Mesotaenium endlicherianum* (Wang et al., 2020)*, Spirogloea muscicola* (Cheng *et al*., 2019)*, Chara braunii* (Nishiyama *et al*., 2018)*, Klebsormidium nitens* (Hori *et al*., 2014)*, Chlorokybus atmophyticus* (Wang et al., 2020)*, Mesostigma viride* (Liang *et al*., 2020)). Protein sequences were downloaded and BLAST searched with the same set of query sequences as above using a local copy of BLASP program (Camacho *et al*., 2009). Further BLAST searches were made in NCBI reference sequences (RefSeq) database (Pruitt *et al*., 2007) and 1kp plant transcriptome database (Leebens-Mack *et al*., 2019). These searches were performed at the CNGB server, https://db.cngb.org/blast/blast/blastp/. For keyword searches, PFMAM domain IDs, PF03552 (Cellulose_synt), PF00535 (Glycos_transf_2), PF13632 (Glyco_trans_2_3), PF14569 (zf-UDP) were searched at phytozome and uniport database servers.

### Sequence analysis

All sequences were retrieved and stored in Microsoft Excel. Simple analyses like identifying the starting amino acid, determining the protein length and searching for presence of catalytic motifs, DD, DxD, ED, QxxRW and FxVTxK were performed within Excel itself. Transmembrane helix (TMH) prediction was performed at phobius server (https://phobius.sbc.su.se/). TMH clustered were determined in Excel. If more than one TMH clusters were present within 100 amino acids of each other, they were deemed to belong to a TMH cluster. A TMH architecture was defined by number of clusters and number of TMHs in each cluster. All sequences were analysed for presence of PFAM domains at motif server (https://www.genome.jp/tools/motif/). All PFAM domains identified were placed in one of 5 main domain groups, GT2, ZB (zinc binding), pilZ, GH (glycosyl hydrolase) and other. A domain group architecture was defined by presence or absence of these domain groups in a given sequence. A domain architecture was also determined for each sequence, where the best domain (based on e value and score) from each of the 5 groups was used.

### Phylogenetics

For producing the guide tree, selected 44210 sequences were aligned using FAMSA (Deorowicz *et al*., 2016) and the guide tree inferred. For classification of CESA sequences, a smaller sub-set of 6580 sequences were aligned with MUSCLE (Edgar, 2004) and a maximum-likelihood tree was produced using FastTree (Price *et al*., 2010).

### Plant material

Secondary cell wall CESA T-DNA mutants, *cesa4^irx5–4^*(SALK_084627), *cesa7^irx3^*^-*7*^ (GABI_819B03), *cesa7^irx3^*^-6^ (SAIL_24_B10) and *cesa8^irx1^*^-*7*^ (GABI_339E12) were obtained from NASC and homozygous plants were identified. Double and triple mutant combinations, *cesa7 cesa8* (*cesa7^irx3^*^-6^ *cesa8^irx1^*^-*7*^), *cesa8 cesa4* (*cesa8^irx1-7^ cesa4^irx5-4^*), *cesa7 cesa4* (*cesa7^irx3^*^-6^ *cesa4^irx5-4^*), *cesa7 cesa8 cesa4* (*cesa7^irx3^*^-6^ *cesa8^irx1^*^-*7*^ *cesa4^irx5-4^*) were obtained by crossing the single mutans. All these mutants have been previously described in Kumar and Turner (2015b).

### DNA Cloning and plant transformation

Plasmids containing *Physcomitrella* CESA cDNA clones, pdp10281 (PpCESA3), pdp21409 (PpCESA4), pdp24095 (PpCESA5), pdp16421 (PpCesA6), pdp34523 (PpCESA7) and pdp39044 (PpCESA8) were obtained from Riken Bioresource Research centre, Japan. These clones were used as template to re-clone the genes in a Gateway compatible vector, pDONOR/pZeo by adding attB adapter sequences in the cloning primers (Table S1). When compared to the CDS sequences from Physcomitrella genome sequencing project, cloned PpCESAs contained some sequence differences. These included 1 bp deletion and a splicing error in PpCESA3 (Pp3c8_7420V3.1), two substitution mutations, P490L and T858A in PpCESA5 (Pp3c2_13330V3.1), two substitution mutations, D344G and R460G in PpCESA6 (DQ902547.1) and two substitution mutations, K205E and P434S in PpCESA7 (Pp3c15_7150V3.1). These sequences were “repaired” using the overlap extension technique described previously (Atanassov *et al*., 2009, Kumar et al., 2017, Kumar et al., 2018a). The repair primers are listed in table SX. *Selaginella moellendorffii* total RNA was prepared from leaf midriff tissue using Aurum RNA extraction kit (Bio-Rad Laboratories) and double stranded cDNA was made with iScript cDNA synthesis kit (Bio-Rad Laboratories). The cDNA was then used to amplify Selaginella CESAs, SmCESA2 (EFJ38258.1) and SmCESA4 (EFJ38300.1) which were then cloned into pDONOR/pZeo. CDS fragments for SmCESA1 (EFJ35421.1), SmCESA3 (EFJ37830.1), all 4 algal CESAs ACESA_MICDE (ADE44904.1, *Micrasterias denticulata*), ACESA_SPISP (HAOX-2004235, *Spirogyra sp*), ACESA_CYLCU (JOJQ-2007927, *Cylindrocystis cushleckae*), ACESA_MESEN (WDCW-2003349, *Mesotaenium endlicherianum*) and the fern CESA UOMY-2008646 from Osmunda were all synthesized commercially by BioBasic, Canada. These sequences were codon optimised for Arabidopsis and included the attB adapter flanking sequences to facilitate Gateway cloning. All sequenced were cloned into pDONOR/pZEO and verified using Sanger sequencing. CDS and protein sequences for all 15 CESA proteins are provided in Table S2. Once validated, all sequences were transferred to a Gateway destination vector VX83 (pAtCESA7::GW, hygromycin plant selection, based on pCB1300 backbone). To make the double CESA plasmids, complete expression cassette for PpCESA8 (pCESA7::PpCESA8:tNOS) was excised and ligated into AscI site of the plasmids containing PpCESA6 and PpCESA7. Sequenced plasmids were transformed into the Agrobacterium strain, pGV3101, which were then transformed into Arabidopsis plants using the floral dip method (Clough & Bent, 1998).

### Plant growth and analysis

T1 seeds were harvested from dipped Arabidopsis plants and selected on ½ MS plates containing 35 µg/mL hygromycin. After growing for 7 days on plates in an incubator, 8-10 independent lines for each construct were transplanted into a 1:1:5 mixture of perlite, vermiculite and compost. Plants were grown for a further 8 weeks on soil under long day conditions (16h/8h day/night, 22°C/18°C temperature and 80% humidity). Col0 WT and the *cesa* mutants were grown on plates without any selection before being transplanted. Vector-only controls for Col0 WT and *cesa* mutants were included in two of the five experiments and no differences were found in the growth patterns or cellulose content as compared to corresponding genotypes grown on non-selection plates. Plant height measurements were taken when plants were 7 weeks old after which 50 mm pieces from the primary inflorescence stem starting at 5 mm above the base, were harvested and stored in 70% ethanol for analysis of cellulose content as described (Kumar & Turner, 2015b). T2 seeds were collected from the secondary inflorescences that were left intact.

Plant height (cm) and cellulose content (% cell wall) were converted into plant height (% complementation) and cellulose content (% complementation) to assess the level of complementation. For statistical analysis, the data was imported into IBM SPSS statistics programme and a univariate analysis of variance with a LSD post hoc test was used to calculate the significance levels for the differences in the means.

### ssNMR

Solid-state NMR analysis was performed as described previously (Kumar et al., 2018a). Briefly, for each genotype, 27 T2 plants were grown on plates and compost as described above. Stem material was harvested when plants were 9 weeks old and stripped of their leaves and siliques. Stems were freeze dried for 72 hours and ground into a fine powder using a bead-mill (TissueLyser II fro Qiagen, UK). The alcohol insoluble residue (AIR) was prepared by extracting twice with 70% ethanol (at 70°C) and once with 1:1 mixture of chloroform:methanol. Samples were then de-starched using amylase and pullulanase.

Up to 50 mg of de-starched AIR powder was analysed by ssNMR in CP/MAS experiments. Carbon-13 magic-angle spinning measurements were carried out at 100.63 MHz using a Bruker Avance III HD spectrometer and 4 mm (rotor o.d.) probe. Spectra were acquired at a spin rate of 10 kHz. Cross-polarisation (CP) spectra were recorded with TOSS spinning sideband suppression, 1 ms contact time and with a recycle delay of 2 s. Carbon spectral referencing is relative to neat tetramethylsilane, carried out by setting the high-frequency signal from an external sample of adamantane to 38.5 ppm. Recorded spectra were exported to Microsoft Excel where they were normalised to make the total signal for each spectrum 100%. Further plotting of the spectra was performed in Excel.

### Digestibility assays

Enzymatic saccharification analysis was performed using the automated system as previously described (Gomez *et al*., 2010). Samples were processed in 96-well plates. For each genotype, up to 24 replicates (6 biological x 4 technical), each weighing 4 mg, were analysed. Enzymatic saccharification and reducing sugars determinations were performed in a liquid handling robotic platform (Tecan Evo 200; Tecan Group Ltd. Männedorf, Switzerland), with the enzyme cocktail Cellic® Ctec3 (7FPU/g) at 50 °C in 25 mmol/L sodium acetate buffer pH 4.5 for 18 h. Automated determination of the reducing sugars released after hydrolysis was performed by colorimetric assay using 3-methyl-2-benzothiazolinone hydrazone (MBTH). Data was expressed as sugar release (nmol/mg).

## Acknowledgements

We would like to thank Karolina Baker who helped generate some of the lines described in this paper. We would also like to thank Rachael Hallam for her assistance in the determination of saccharification. The work was funded by BBSRC grant BB/X016919/1 and The Leverhulme Trust grant RPG-2020-257. The authors declare no conflict of interest.

## Author Contributions

MK and ST designed the research and wrote the paper. MK and LDG performed research and analyzed data.

## Accession numbers

Accession numbers for Arabidopsis genes mentioned in this paper are AT5G44030 (CESA04), AT5G17420 (CESA07), AT4G18780 (CESA08). Accession numbers for 15 heterologous proteins are provided in Table S2.

## References

Atanassov, I., Atanassov, I., Etchells, J. P. and Turner, S. (2009) A simple, flexible and efficient PCR-fusion/Gateway cloning procedure for gene fusion, site-directed mutagenesis, short sequence insertion and domain deletions and swaps. Plant Methods, 5, 14.

Blum, M., Gamper, H. A., Waldner, M., Sierotzki, H. and Gisi, U. (2012) The cellulose synthase 3 (CesA3) gene of oomycetes: structure, phylogeny and influence on sensitivity to carboxylic acid amide (CAA) fungicides. Fungal Biology, 116, 529–542.

Brawley, S. H., Blouin, N. A., Ficko-Blean, E., Wheeler, G. L., Lohr, M., Goodson, H. V., et al. (2017) Insights into the red algae and eukaryotic evolution from the genome of Porphyra umbilicalis (Bangiophyceae, Rhodophyta). Proc. Natl. Acad. Sci. USA, 114, E6361–E6370.

Camacho, C., Coulouris, G., Avagyan, V., Ma, N., Papadopoulos, J., Bealer, K., et al. (2009) BLAST plus: architecture and applications. BMC Bioinformatics, 10.

Chan, W. S., Kwok, A. C. M. and Wong, J. T. Y. (2019) Knockdown of Dinoflagellate Cellulose Synthase CesA1 Resulted in Malformed Intracellular Cellulosic Thecal Plates and Severely Impeded Cyst-to-Swarmer Transition. Frontiers in Microbiology, 10.

Chen, S. L., Ehrhardt, D. W. and Somerville, C. R. (2010) Mutations of cellulose synthase (CESA1) phosphorylation sites modulate anisotropic cell expansion and bidirectional mobility of cellulose synthase. Proc. Natl. Acad. Sci. USA, 107, 17188–17193.

Cheng, S. F., Xian, W. F., Fu, Y., Marin, B., Keller, J., Wu, T., et al. (2019) Genomes of Subaerial Zygnematophyceae Provide Insights into Land Plant Evolution. Cell, 179, 1057-+.

Clough, S. J. and Bent, A. F. (1998) Floral dip: a simplified method for Agrobacterium-mediated transformation of Arabidopsis thaliana. Plant J., 16, 735–743.

Cosgrove, D. J. (2024) Structure and growth of plant cell walls. Nature Reviews Molecular Cell Biology, 25, 162–167.

de Vries, L., Guevara-Rozo, S., Cho, M., Liu, L. Y., Renneckar, S. and Mansfield, S. D. (2021) Tailoring renewable materials via plant biotechnology. Biotechnology for Biofuels, 14.

Deorowicz, S., Debudaj-Grabysz, A. and Gudys, A. (2016) FAMSA: Fast and accurate multiple sequence alignment of huge protein families. Scientific Reports, 6.

Ding, Y., Pang, Z. Q., Lan, K., Yao, Y., Panzarasa, G., Xu, L., et al. (2022) Emerging Engineered Wood for Building Applications. Chemical Reviews.

Dupree, R., Simmons, T. J., Mortimer, J. C., Patel, D., Iuga, D., Brown, S. P., et al. (2015) Probing the Molecular Architecture of Arabidopsis thaliana Secondary Cell Walls Using Two- and Three-Dimensional C-13 Solid State Nuclear Magnetic Resonance Spectroscopy. Biochemistry, 54, 2335–2345.

Edgar, R. C. (2004) MUSCLE: a multiple sequence alignment method with reduced time and space complexity. BMC Bioinformatics, 5, 1–19.

Fang, Y. H., Qin, X., Liao, Q. G., Du, R., Luo, X. Z., Zhou, Q., et al. (2021) The genome of homosporous maidenhair fern sheds light on the euphyllophyte evolution and defences. Nature Plants, 8, 1024-+.

Fernandes, A. N., Thomas, L. H., Altaner, C. M., Callow, P., Forsyth, V. T., Apperley, D. C., et al. (2011) Nanostructure of cellulose microfibrils in spruce wood. Proc. Natl. Acad. Sci. USA, 108, E1195–E1203.

Gomez, L. D., Whitehead, C., Barakate, A., Halpin, C. and McQueen-Mason, S. J. (2010) Automated saccharification assay for determination of digestibility in plant materials. Biotechnology for Biofuels, 3.

Goodstein, D. M., Shu, S. Q., Howson, R., Neupane, R., Hayes, R. D., Fazo, J., et al. (2012) Phytozome: a comparative platform for green plant genomics. Nucleic Acids Res., 40, D1178–D1186.

Harris, D. M., Corbin, K., Wang, T., Gutierrez, R., Bertolo, A. L., Petti, C., et al. (2012) Cellulose microfibril crystallinity is reduced by mutating C-terminal transmembrane region residues CESA1(A903V) and CESA3(T942I) of cellulose synthase. Proc. Natl. Acad. Sci. USA, 109, 4098–4103.

Hori, K., Maruyama, F., Fujisawa, T., Togashi, T., Yamamoto, N., Seo, M., et al. (2014) Klebsormidium flaccidum genome reveals primary factors for plant terrestrial adaptation. Nature Communications, 5.

Jia, C., Chen, C. J., Mi, R. Y., Li, T., Dai, J. Q., Yang, Z., et al. (2019) Clear Wood toward High-Performance Building Materials. Acs Nano, 13, 9993–10001.

Kumar, M., Atanassov, I. and Turner, S. (2017) Functional Analysis of Cellulose Synthase (CESA) Protein Class Specificity. Plant Physiol., 173, 970–983.

Kumar, M., Mishra, L., Carr, P., Pilling, M., Gardner, P., Mansfield, S. D., et al. (2018a) Exploiting CELLULOSE SYNTHASE (CESA) Class Specificity to Probe Cellulose Microfibril Biosynthesis. Plant Physiol., 177, 151–167.

Kumar, M., Mishra, L., Carr, P., Pilling, M., Gardner, P., Mansfield, S. D., et al. (2018b) Exploiting CELLULOSE SYNTHASE (CESA) class-specificity to probe cellulose microfibril biosynthesis. Plant Physiol., 177, 151–167.

Kumar, M. and Turner, S. (2015a) Plant cellulose synthesis: CESA proteins crossing kingdoms. Phytochemistry, 112, 91–99.

Kumar, M. and Turner, S. (2015b) Protocol: a medium-throughput method for determination of cellulose content from single stem pieces of Arabidopsis thaliana. Plant Methods, 11, 46.

Kumar, M., Wightman, R., Atanassov, I., Gupta, A., Hurst, C. H., Hemsley, P. A., et al. (2016) S-Acylation of the cellulose synthase complex is essential for its plasma membrane localization. Science, 353, 166–169.

Leebens-Mack, J. H., Barker, M. S., Carpenter, E. J., Deyholos, M. K., Gitzendanner, M. A., Graham, S. W., et al. (2019) One thousand plant transcriptomes and the phylogenomics of green plants. Nature, 574, 679-+.

Li, F. W., Brouwer, P., Carretero-Paulet, L., Cheng, S. F., de Vries, J., Delaux, P. M., et al. (2018) Fern genomes elucidate land plant evolution and cyanobacterial symbioses. Nature Plants, 4, 460–472.

Li, F. W., Nishiyama, T., Waller, M., Frangedakis, E., Keller, J., Li, Z., et al. (2020) Anthoceros genomes illuminate the origin of land plants and the unique biology of hornworts. Nature Plants, 6, 259–272.

Li, X. X., Speicher, T. L., Dees, D. C. T., Mansoori, N., McManus, J. B., Tien, M., et al. (2019) Convergent evolution of hetero-oligomeric cellulose synthesis complexes in mosses and seed plants. Plant J., 99, 862–876.

Liang, Z., Geng, Y. K., Ji, C. M., Du, H., Wong, C. E., Zhang, Q., et al. (2020) Mesostigma viride Genome and Transcriptome Provide Insights into the Origin and Evolution of Streptophyta. Advanced Science, 7.

Little, A., Schwerdt, J. G., Shirley, N. J., Khor, S. F., Neumann, K., O’Donovan, L. A., et al. (2018) Revised Phylogeny of the Cellulose Synthase Gene Superfamily: Insights into Cell Wall Evolution. Plant Physiol., 177, 1124–1141.

Marchant, D. B., Sessa, E. B., Wolf, P. G., Heo, K., Barbazuk, W. B., Soltis, P. S., et al. (2019) The C-Fern (Ceratopteris richardit) genome: insights into plant genome evolution with the first partial homosporous fern genome assembly. Scientific Reports, 9.

Mergner, J., Frejno, M., List, M., Papacek, M., Chen, X., Chaudhary, A., et al. (2020) Mass-spectrometry-based draft of the Arabidopsis proteome. Nature, 579, 409–414.

Niklas, K. J. (2004) The cell walls that bind the tree of life. Bioscience, 54, 831–841.

Nishiyama, T., Sakayama, H., de Vries, J., Buschmann, H., Saint-Marcoux, D., Ullrich, K. K., et al. (2018) The Chara Genome: Secondary Complexity and Implications for Plant Terrestrialization. Cell, 174, 448-+.

Norris, J. H., Li, X., Huang, S., van de Meene, A., Tran, M. L., Killeavy, E., et al. (2017a) Functional specialization of cellulose synthase isoforms in a moss shows parallels with seed plants. Plant Physiol.

Norris, J. H., Li, X. X., Huang, S. X., Van de Meene, A. M. L., Tran, M. L., Killeavy, E., et al. (2017b) Functional Specialization of Cellulose Synthase Isoforms in a Moss Shows Parallels with Seed Plants. Plant Physiol., 175, 210–222.

Pancaldi, F., van Loo, E. N., Schranz, M. E. and Trindade, L. M. (2022) Genomic Architecture and Evolution of the Cellulose synthase Gene Superfamily as Revealed by Phylogenomic Analysis. Front. Plant. Sci., 13.

Pang, Z. L., McKee, L. S., Srivastava, V., Klinter, S., Diaz-Moreno, S. M., Orlean, P., et al. (2020) Analysis of a cellulose synthase catalytic subunit from the oomycete pathogen of cropsPhytophthora capsici. Cellulose, 27, 8551–8565.

Price, M. N., Dehal, P. S. and Arkin, A. P. (2010) FastTree 2-Approximately Maximum-Likelihood Trees for Large Alignments. PLoS One, 5.

Pruitt, K. D., Tatusova, T. and Maglott, D. R. (2007) NCBI reference sequences (RefSeq): a curated non-redundant sequence database of genomes, transcripts and proteins. Nucleic Acids Res., 35, D61–D65.

Purushotham, P., Ho, R. and Zimmer, J. (2020) Architecture of a catalytically active homotrimeric plant cellulose synthase complex. Science, eabb2978.

Scavuzzo-Duggan, T. R., Chaves, A. M., Singh, A., Sethaphong, L., Slabaugh, E., Yingling, Y. G., et al. (2018) Cellulose synthase ‘class specific regions’ are intrinsically disordered and functionally undifferentiated. Journal of integrative plant biology.

Schubert, M., Panzarasa, G. and Burgert, I. (2022) Sustainability in Wood Products: A New Perspective for Handling Natural Diversity. Chemical Reviews.

Sena, J. S., Lachance, D., Duval, I., Nguyen, T. T. A., Stewart, D., Mackay, J., et al. (2019) Functional Analysis of the PgCesA3 White Spruce Cellulose Synthase Gene Promoter in Secondary Xylem. Front. Plant. Sci., 10.

Song, J. W., Chen, C. J., Zhu, S. Z., Zhu, M. W., Dai, J. Q., Ray, U., et al. (2018) Processing bulk natural wood into a high-performance structural material. Nature, 554, 224-+.

Tran, M. L., McCarthy, T. W., Sun, H., Wu, S. Z., Norris, J. H., Bezanilla, M., et al. (2018) Direct observation of the effects of cellulose synthesis inhibitors using live cell imaging of Cellulose Synthase (CESA) in Physcomitrella patens. Scientific Reports, 8.

Tsekos, I. (1999) The sites of cellulose synthesis in algae: Diversity and evolution of cellulose-synthesizing enzyme complexes. J. Phycol., 35, 635–655.

Updegraff, D. M. (1969) Semimicro determination of cellulose in biological materials. Anal. Biochem., 32, 420-&.

Wan, T., Liu, Z. M., Li, L. F., Leitch, A. R., Leitch, I. J., Lohaus, R., et al. (2018) A genome for gnetophytes and early evolution of seed plants. Nature Plants, 4, 82–89.

Wang, S. B., Li, L. Z., Li, H. Y., Sahu, S. K., Wang, H. L., Xu, Y., et al. (2020) Genomes of early-diverging streptophyte algae shed light on plant terrestrialization. Nature Plants, 6, 95-+.

Xiao, S. L., Chen, C. J., Xia, Q. Q., Liu, Y., Yao, Y., Chen, Q. Y., et al. (2021) Lightweight, strong, moldable wood via cell wall engineering as a sustainable structural material. Science, 374, 465–471.

Xue, J., Purushotham, P., Acheson, J. F., Ho, R. Y., Zimmer, J., McFarlane, C., et al. (2022) Functional characterization of a cellulose synthase, CtCESA1, from the marine red alga *Calliarthron tuberculosum* (Corallinales). J. Exp. Bot., 73, 680-695.

Zhang, J., Fu, X. X., Li, R. Q., Zhao, X., Liu, Y., Li, M. H., et al. (2020) The hornwort genome and early land plant evolution. Nature Plants, 6, 107–118.

